# Deciphering molecular determinants of GPCR-G protein receptor interactions by complementary integrative structural biology methods

**DOI:** 10.1101/2024.10.21.619395

**Authors:** Jérôme Castel, Thomas Botzanowski, Ieva Brooks, Alexandre Frechard, Gilbert Bey, Marine Schroeter, Elise Del Nero, François Debaene, Fabrice Ciesielski, Denis Zeyer, Sarah Cianferani, Renaud Morales

## Abstract

Many physiological processes are dependent on G protein-coupled receptors (GPCRs), the biggest family of human membrane proteins and a significant class of therapeutic targets. Once activated by external stimuli, GPCRs use G proteins and arrestins as transducers to generate second messengers and trigger downstream signaling, leading to diverse signaling profiles. The G protein-coupled bile acid receptor 1 (GPBAR1, also known as Takeda G protein-coupled receptor 5, TGR5) is a class A bile acid membrane receptor that regulates energy homeostasis and glucose and lipid metabolism. GPBAR1/G protein interactions are implicated in the prevention of diabetes and the reduction of inflammatory responses, making GPBAR1 a potential therapeutic target for metabolic disorders. Here, we present an integrated structural biology approach combining hydrogen/deuterium exchange mass spectrometry (HDX-MS) and cryo-electron microscopy (cryo-EM) to identify the molecular determinants of GPBAR1 conformational dynamics upon G protein binding. Thanks to extensive optimization of both HDX-MS and cryo-EM workflows, we achieved over 99% sequence coverage along with a 2.5-Å resolution structure, both of which are state-of-the-art and solely obtained for complete GPCR complexes. Altogether, our results provide information on the under-investigated GPBAR1 binding mode to its cognate G protein, pinpointing the synergic and powerful combination of higher (cryo-EM) and lower (HDX-MS) resolution structural biology techniques to dissect GPCR/G protein binding characteristics.

**Short statement:** This work highlights the utility of integrating cryo-EM and HDX-MS for studying large multiprotein complexes such as GPCR/G protein complexes. Cryo-EM offers high-resolution structural details, while HDX-MS reveals the dynamic conformational changes during assembly, providing a comprehensive structural view of difficult-to-study membrane protein systems.

## 1 INTRODUCTION

G protein-coupled receptors (GPCRs) constitute the largest and the most diverse family of cell-surface receptors, with around 800 identified in humans. They play an essential role in the regulation of physiological functions (such as vision, olfactory perception, neurotransmission, pain, immunity…), and their dysfunctions or mutations are associated with a variety of diseases (Alzheimer’s disease, hypertension, bronchial asthma, multiple sclerosis, and diabetes, on so on). GPCRs are, therefore, widely recognized as important therapeutic targets for pharmaceutical development, with 30-40% of all drugs approved by the US Food and Drug Administration (FDA) known to specifically target GPCRs. From a structural point of view, all GPCRs share a common architecture composed of seven transmembrane α-helices (TM1-TM7) connected by three extracellular loops (ECLs) and three intracellular loops (ICLs). GPCRs have an extracellular N-terminus that can participate in ligand binding, especially in class B and C GPCRs, while the intracellular C-terminus interacts with various intracellular signaling proteins, including G proteins, kinases, and arrestins.

GPCR activation is induced by extracellular signals, such as photons, ligands, or hormones, allowing them to interact with signaling molecules[1–3], among which the G proteins are the best characterized[4]. These are heterotrimeric complexes composed of alpha (Gα), beta (Gβ), and gamma (Gγ) subunits. G proteins can be distinguished by their Gα subunits, which are grouped into four families based on sequence similarities and functional results (Gαs, Gαi/o, Gαq/11, and Gα12/13)[5]. In the basal (resting) state, the Gα subunit, whose catalytic site is occupied by a guanosine diphosphate (GDP) nucleotide, interacts with the Gβ and Gγ subunits. Activation of G proteins and initiation of signaling cascades are achieved by nucleotide dissociation from the Gα subunit. The interaction of a GPCR with a G protein triggers the exchange of GDP into guanosine triphosphate (GTP), followed by dissociation of the Gα subunit first and the Gβ/Gγ subunits second. The GTP-bound Gα subunit or Gβ/γ subunits then interact with and regulate effector proteins (ion channels, phospholipases…)[6, 7]. Although some GPCRs interact with a single G protein, many can bind to one or more Gα subunits, inducing the activation of several effector proteins[8, 9].

The structural characterization of GPCR/G-protein (GPCR/G) interactions enables understanding of GPCR signaling mechanisms and the development of new drugs targeting these receptors. X-ray crystallography, cryo-EM[10, 11], and NMR have been the predominant methodologies deployed to obtain high-resolution 3D structures and the study of GPCRs complexed to ligands and G proteins. The first structure of a GPCR/G protein complex was obtained by crystallography in 2011, revealing β2A adrenergic receptor interacting with a heterotrimeric Gs protein in a nucleotide-free state and stabilized by a nanobody (Nb35)[12]. Given the challenges in stabilizing GPCR/G complexes in an active conformation, several strategies have been developed. Firstly, apyrase was used to trap the GPCR/G complex in a nucleotide-free state[11, 13]. Secondly, a single-domain antibody (nanobody) was added to stabilize the interface between the Gβ/Gγ and GD subunits within the protein G, thus stabilizing the GPCR/G complex[14, 15]. Subsequently, technological breakthroughs in cryo-EM have enabled the resolution of numerous GPCR/G structures with the Gs subfamily (CTR[16], GLP1[17], etc.), Gi (opsin[18], MOR[19], endothelin-1-ETB[20]) or Go (5HT1BR[21]). However, most of these structures are static snapshots of the most stable state and do not provide complete visualization of the conformational states and dynamics of GPCRs. Indeed, most of the transient and metastable states present in solution are often not “captured” in crystallography or cryo-EM. Over the past two decades, MS has established itself as a powerful, complementary approach to the more conventional high-resolution biophysical techniques used in structural biology to probe the structural architecture and dynamics of GPCRs and their related complexes[22–24]. In particular, MS has been used to understand the ability of different molecules or ligands to modulate the structural dynamics of GPCRs, including during the formation of complexes between G proteins and receptors[25–27]. The HDX-MS approach provides valuable and essential information on the dynamics and mechanisms leading to complex formation between GPCRs and GPCR/G complexes[28–30]. HDX-MS enables the monitoring of solvent exchange by exposing the hydrogens within a protein molecule to a solvent containing the heavy hydrogen isotope—deuterium. HDX-MS is a powerful tool for studying protein structures, stability, folding, conformational dynamics, and binding regions. Since the first use of HDX-MS on a GPCR/G complex involving β2A[31] in 2011, several additional HDX-MS studies have been carried out on various GPCRs. These studies have led to a better understanding of the structural mechanism linking complex formation and GDP nucleotide release[31], as well as highlighting the existence of transient interactions during the coupling process[32, 33].

Here, we used full-length active GPBAR1 GPCR bound to its agonist P395 ligand in interaction with heterotrimeric Gs (comprising Gαs, Gβ, and Gγ subunits; and stabilized by Nb35) to illustrate the synergy and complementarity of HDX-MS and cryo-EM integration in investigating the molecular determinants and interaction dynamics of GPCR/G protein. GPBAR1 is a class A bile acid membrane receptor that regulates energy homeostasis and glucose and lipid metabolism. In particular, GPBAR1/Gs interactions are involved in the prevention of diabetes and the reduction of inflammatory responses, making GPBAR1 a potential therapeutic target for several diseases, such as obesity or atherosclerosis. We present a high-resolution 2.5-Å cryo-EM structure of the GPBAR1/G complex, along with a comprehensive HDX-MS derived overview of the conformational dynamics while interacting with its Gs subunits of this under-investigated GPCR.

## 2 RESULTS

### 2.1 Purification and intact mass analysis of the GPBAR1/G complex

To form the active GPBAR1/G complex, GPBAR1 was co-expressed with Gαs, Gβ, and Gγ in Sf21 insect cells and purified in the presence of P395, a highly potent synthetic agonist (see Materials and Methods for details). Further stabilization of the complex was achieved using camelid antibody, Nb35, which binds at the Gαs-Gβ interface. To assess the purity and homogeneity of the different protein assemblies, we first performed an intact mass analysis of the constituents using mass photometry (MPhoto), a label-free technique that quantifies the molecular weights of biomolecules at the single-molecule level by interferometric detection of scattered light (Figure 1, and Supplementary data, Figure A)[34]. Figure 1 shows MPhoto experiments of GPBAR1 and the GPBAR1/G complex (Figures 1.B-C). For GPBAR1, MPhoto shows one single peak at 51 ± 16 kDa, in agreement with the theoretical mass (50.8 kDa). A minor species at 174 ± 75 kDa is also detected at higher mass, which could be attributed to GPBAR1 aggregation during sample dilution since the detergent concentration is below the critical micelle concentration (CMC). For the GPBAR1/G complex, MPhoto then revealed the sample homogeneity as a single sharp peak, with a mass of 239 ± 23 kDa, which is higher than the expected mass for the complex (∼158 kDa). This mass difference of +81 kDa is in agreement with the reported size of LMNG micelles (50-80 kDa). Although it cannot be ruled out that this single peak is composed of a mixture of empty LMNG and GPBAR1/G-filled micelles, MPhoto suggests that the 1:1 GPBAR1:G complex is formed. Note that a minor distribution at 50 ± 17 kDa is also detected, which could correspond to an excess of G protein subunits present in the solution.

**Figure 1:**
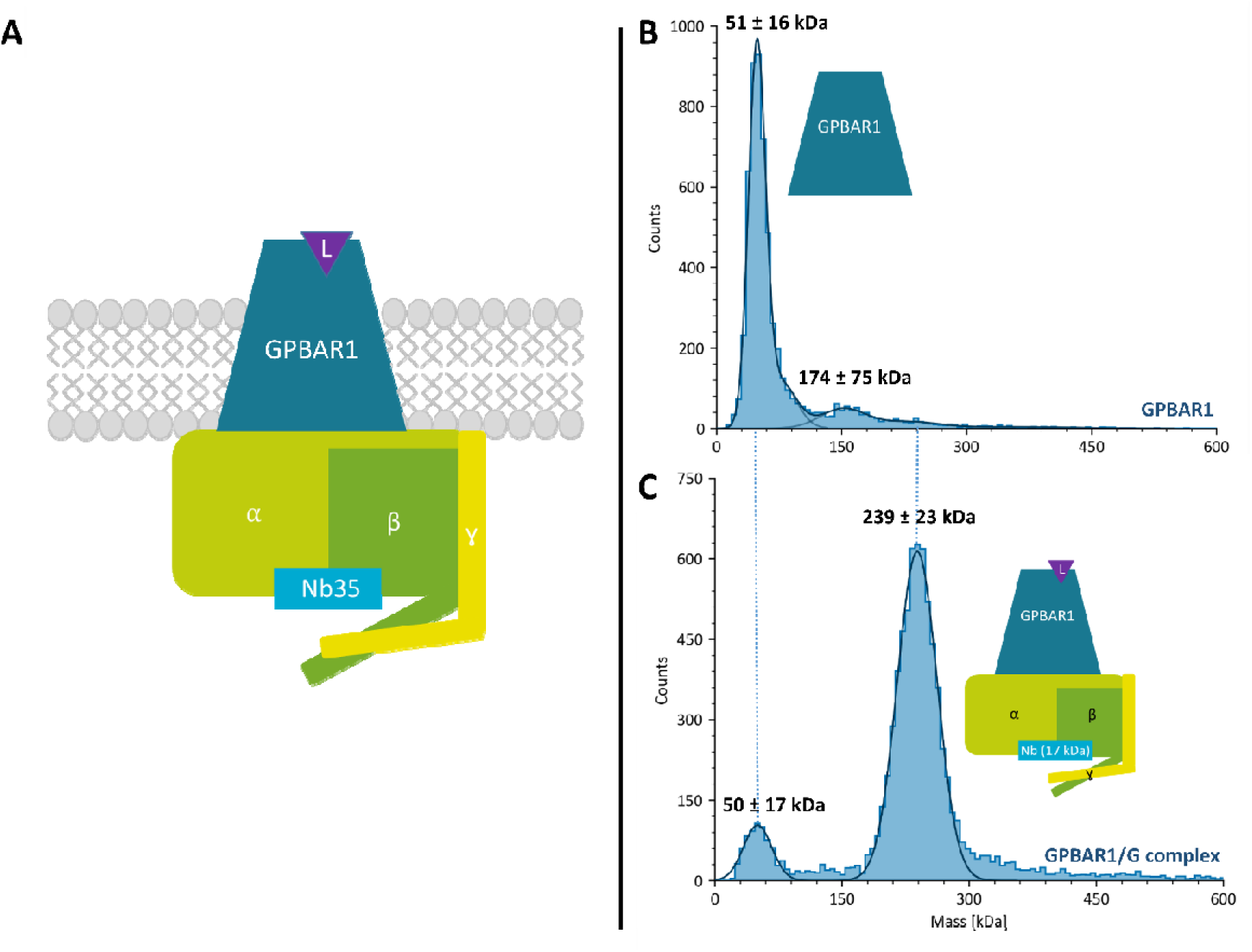
MPhoto analyses of GPBAR1 alone and the GPBAR1/G complex. (A) Schematic representation of the GPBAR1/G complex. The GPBAR1 receptor (51 kDa) is activated by the agonist ligand P395. The different G protein subunits are shown: Gαs in light green (43 kDa), Gβ in dark green (39 kDa), and Gγ (8 kDa) in yellow. The complex is stabilized by the nanobody Nb35 (17 kDa), shown in cyan. (B). Histogram of the mass distribution of the GPBAR1. (C) Histogram of the mass distribution of the GPBAR1/G complex. The continuous curve corresponds to the mass distributions of the majority of species fitted with a Gaussian function.

### 2.2 HDX-MS for conformational dynamics of GPBAR1/Gs complex

As optimization of digestion conditions is of the utmost importance in HDX-MS workflows, we first carried out an exhaustive exploration of all pre-analytical steps of the HDX-MS workflow on GPBAR1 in LMNG, including the amount injected, the nature of the protease and the quenching buffer to achieve the optimum conditions for concomitantly yielding the highest number of peptides (# peptides); sequence coverage (Sc), and redundancy (R) (see supplementary data Figures D and E). HDX-MS optimization gave 175 peptides covering >99% of the GPBAR1 sequence within the GPBAR1/Gs complex, with an average peptide redundancy of 6.7, which is quite remarkable and state-of-the-art for GPCRs. It should be noted that under similar conditions, all G-protein subunits and the Nb35 nanobody gave extensive data (Sc>94%, R>11), enabling us to obtain precise information on conformational changes occurring in all the constituent partners of the GPBAR1/G complex (see supplementary data, Table A).

Overall, GPBAR1 is highly dynamic upon binding to G-protein along its sequence, with almost all regions (including TM2-3 and TM5-7) strongly protected from H/D exchange after complex formation (Figure 2.A). It is worth noting that the peptides in the BRIL sequence showed no difference in D incorporation between the two states (see supplementary data, Figure G), underlining the consistency of the HDX-MS data set.

**Figure 2:**
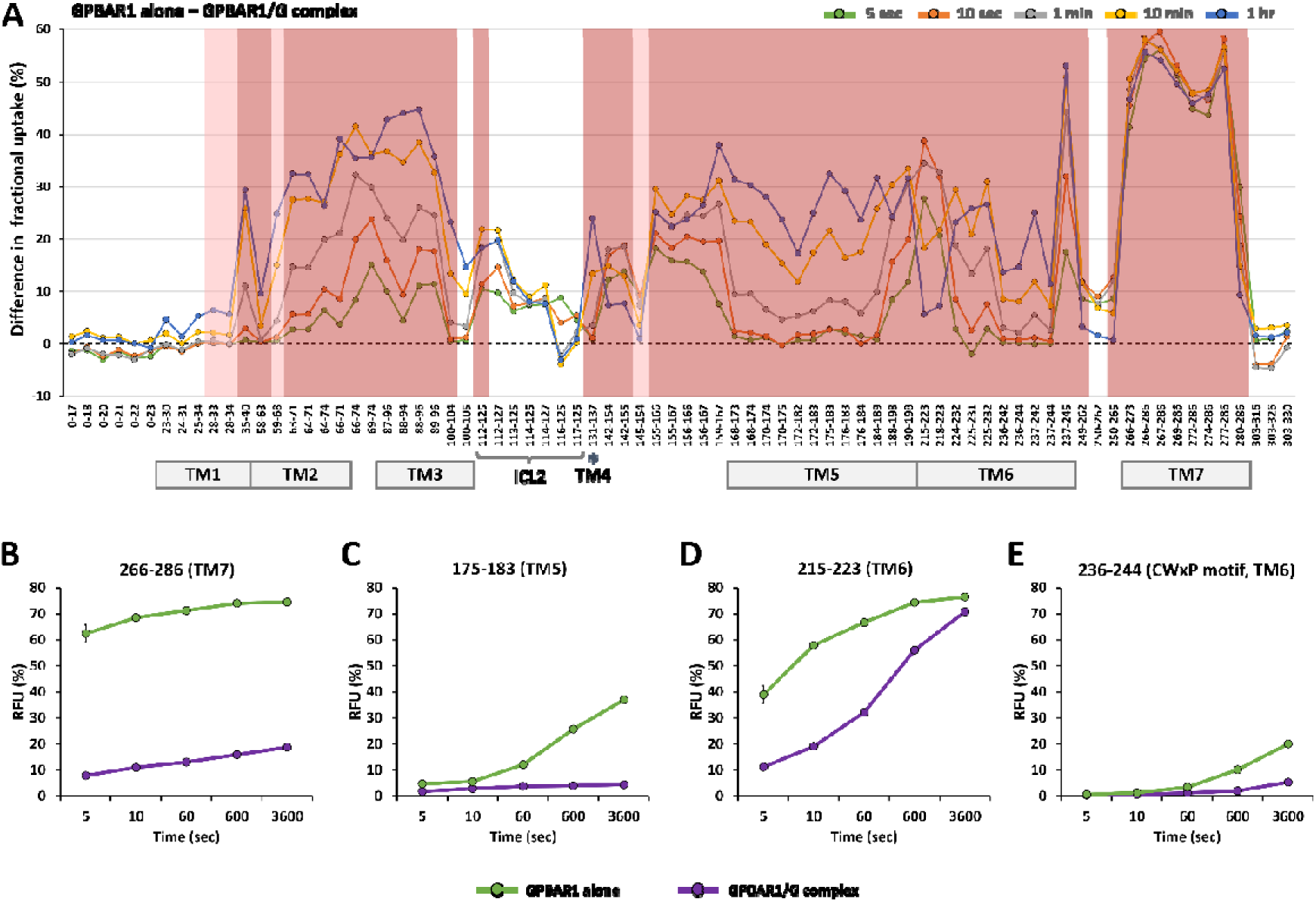
Differences in relative D uptake between GPBAR1 alone and complexed for each peptide identified at different deuteration times (A). Boxed peptides show statistically significant differences in D incorporation (MEMHDX, p-value 0.01), between 5 and 10% (light red) and above 10% (dark red). Relative D incorporation for 4 peptides from GPBAR1 helices TM7 (B), TM5 (C), and TM6 (D, E).

Focusing on the TM domains, the largest increase in D uptake is observed for the TM7 helix, with incorporation differences between 40% and 60% between the two states (Figure 2B). Similarly, the TM5 helix is highly dynamic, reaching a D incorporation difference of almost 35% after one hour of labeling (Figure 2C). These local rearrangements are also accompanied by movements of the intracellular ends of TM5 and TM6 helices. For example, peptide 215-223 from TM6 shows protection against H/D exchange at short time points (5 s to 1 min), whereas no difference is observed at longer time points, suggesting greater exposure of this region to the solvent (Figure 2D). The TM2 and TM3 helices also exhibit a very high level of dynamics during complex formation, with some peptides showing 30 to 45% H/D exchange for the longest labeling points (peptides 63-71, 66-71, 87-96 or 88-96 for example). Another important region for GPCR activation is the CWxP motif of the ligand-binding pocket found in the TM6 (amino acids at positions 236, 237, and 239 in our case). As expected, the CWxP motif (peptide 236-244) has very low solvent accessibility (20% incorporation in D after 1 hour of labeling), in agreement with the fact that the ligand binding pocket is buried in GPBAR1 (Figure 2E). HDX-MS observations confirm the general mechanism of GPCR/G interaction, where Cys236 contributes to the reorganization of TM6-TM7 interactions during receptor activation, while Trp237 movement enables the intracellular end of TM6 to move[35]. Due to their inherent flexibility, the intracellular loops (ICL1-3) are challenging to assess using HDX-MS. In our HDX-MS experiments, only peptides from ICL2 could be reliably monitored, with peptide 112-125 exhibiting significant protection from D uptake upon GPBAR1/Gs complex formation. (Figure 2A).

We then investigated the dynamics of the three Gαs, Gβ, Gγ subunits of the G protein receptor (Figure 3). Significantly different dynamics are observed for the three subunits, with Gαs being the most flexible, Gβ being less affected, and finally Gγ being almost unaffected by GPBAR1 binding. In more detail, Gα shows protected (red) and deprotected (blue) regions.

**Figure 3:**
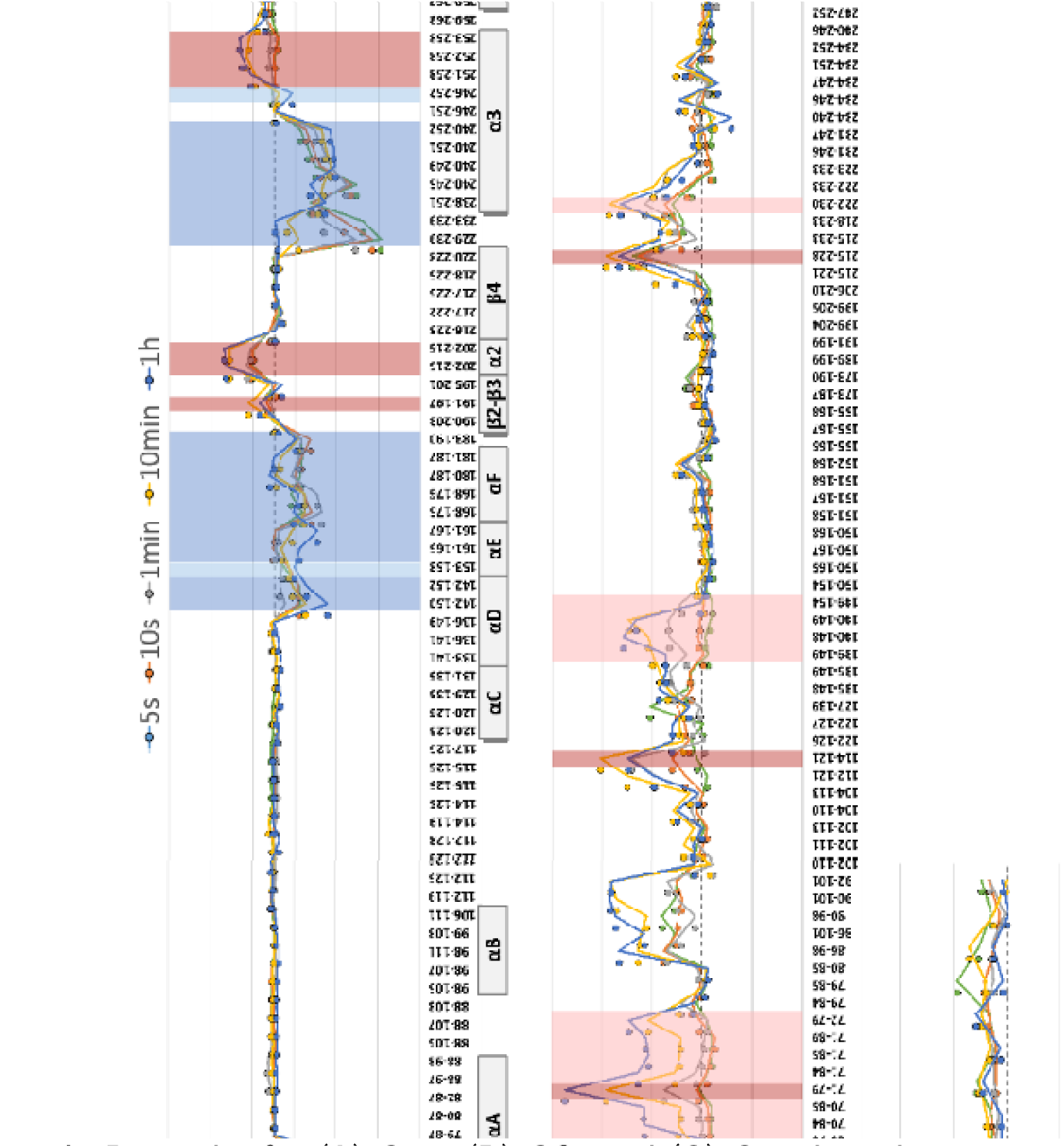
Relative differences in D uptake for (A) Gαs, (B) Gβ, and (C) Gγ when the G protein is alone or in complex with GPBAR1 for each peptide identified at different deuteration times. Boxed peptides show statistically significant differences in D incorporation (MEMHDX, p-value 0.01). Peptides framed in red show a decrease in solvent accessibility after complex formation (protection), while those in blue show an increase (deprotection).

The C-terminal region of Gαs is strongly protected during complex formation, as depicted by the D incorporation profile of peptide 358-370 (end of α5 helix, Figure 4A). After one hour of labeling, this peptide reaches a D incorporation difference of nearly 40%. In contrast to the C-terminal end of the α5 helix, which is strongly protected, the N-terminus is more solvent-exposed. For instance, peptide 347-352 has a deuterium incorporation difference of up to 40%. Moreover, the lack of statistically significant differences for β1 strand, as described in previous studies of class A GPCR (A_2_A, β_2_A)[30, 32], may suggest a more superficial insertion of the α5 helix in GPBARs receptors[36, 37], Therefore, subtle differences in D incorporation at this interaction interface could not be visualized in HDX-MS.

**Figure 4:**
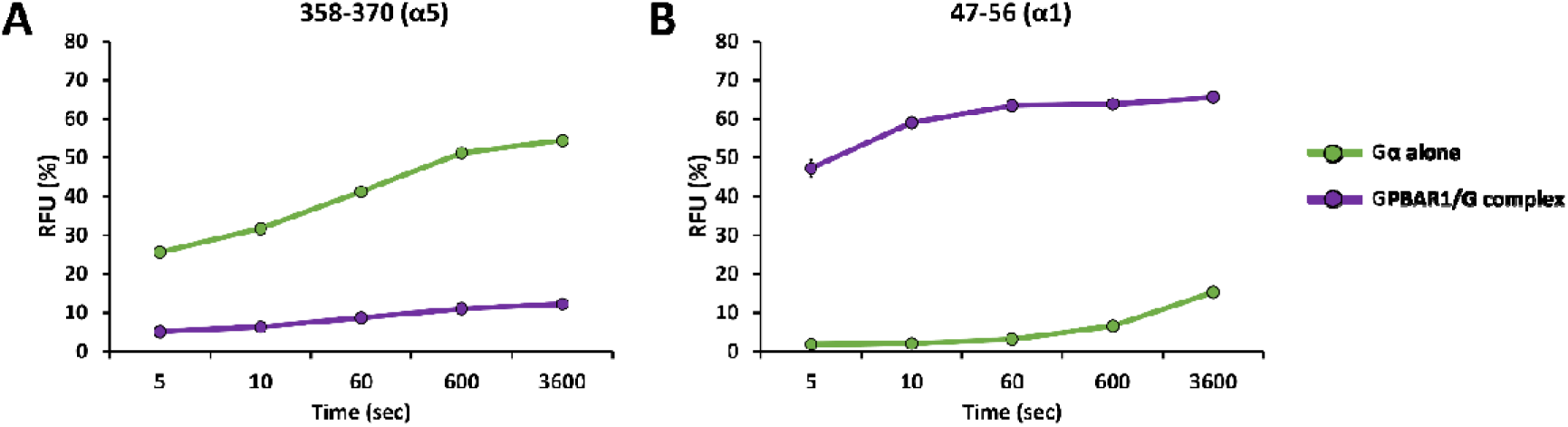
Relative incorporation in D for 2 peptides from Gα digestion obtained in HDX-MS.

Reduced accessibility upon deuteration is observed in the GPBAR/Gs complex for αN helix (peptides 11-18 and 19-25), β2-β3 strands (peptide 191-197), α4 helix (peptide 322-328), and β6 strand (peptide 327-339), consistent with the cryo-EM structure[36] and their involvement in the complex formation.

Overall, peptides of the helical domains of Gαs (helices αA to αD, respectively) show no significant differences in D incorporation, except for peptides 68-78 and 142-150 which exhibit increased solvent exposure following GPBAR1 binding (Figure 3).

Furthermore, in the GPBAR/Gs complex, certain regions of the Gαs GTPase domain become more exposed to the solvent. Helices α1 (e.g. peptide 47-56, Figure 4B), as well as regions of α3, αG, and β5, exhibit significantly increased D incorporation after complex formation (50-60% incorporation difference for peptide 47-56). Solvent exposure of the region formed by the loop connecting β1 to α1 and α1 shows that this region is particularly flexible and dynamic during complex formation, irrespective of the GPCR family. Other regions of the GTPase domain[36], such as αN, α2, and α5 helices, regions of α3, α4, β2, β3, and β6 strands), are protected, consistent with the Gα subunit interacting with GPBAR1.

Overall, our HDX-MS results are consistent with previous studies showing that the helical domain rotated by 127° to allow interaction between the receptor and the rest of the G protein[12].

Conversely, the Gβ subunit is significantly less affected by GPBAR1 binding, with fewer regions exhibiting protection from D incorporation. Lower D incorporation for peptides 71-79, 114-121 and 215-228 is observed during GPBAR1/Gs complex formation. For peptide 215-228 (Figure 5A), a significant difference in incorporation and dynamics can be observed between the two states, whereas peptide 71-79 (Figure 5B) mainly shows only a significant difference in dynamics. Finally, as expected, Gγ shows no statistically relevant region. This result is not surprising, as this small subunit (8 kDa) is located at the periphery of the complex and the conformational changes linked to the interaction between GPBAR1 and Gα are too distant to be visible on Gγ.

**Figure 5:**
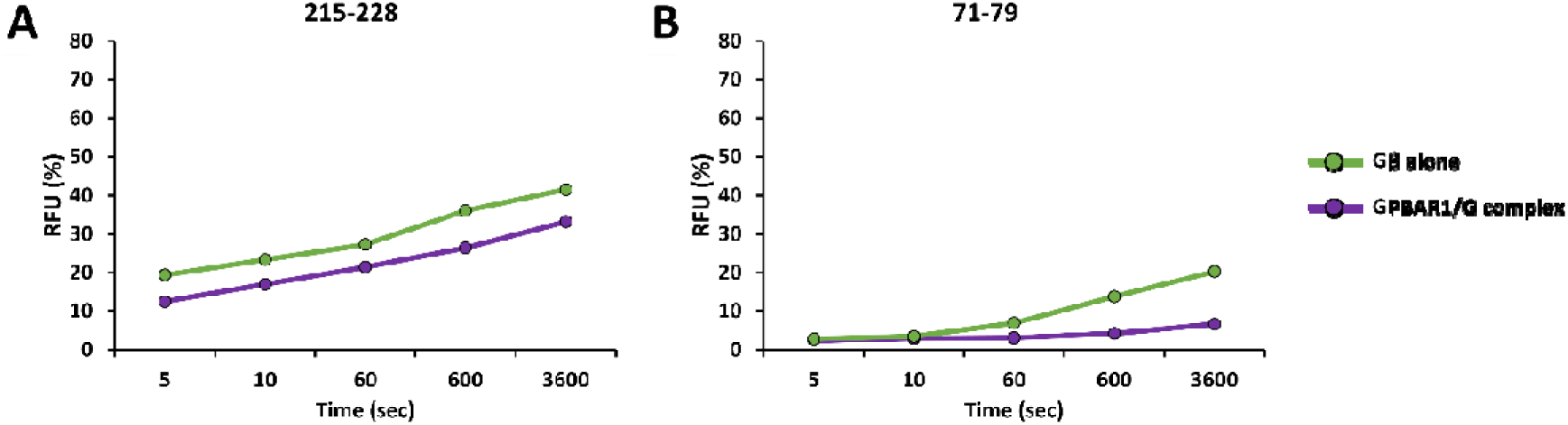
Relative incorporation in D for 2 peptides from Gβ digestion obtained in HDX-MS.

### 2.3 Cryo-EM structure of the GPBAR1/G complex

We obtained two cryo-EM structures of the GPBAR-Gs complex bound to the P395 ligand. The first one, referred to as the core structure, has achieved the highest average resolution of 2.5 Å (Figure 6A, C), showing all the stable parts of the structure, including as well as the P395 ligand. The second structure, with an average resolution of 2.9 Å (Figure 6B, D), is very similar to the core structure but additionally reveals the flexible, weakly resolved α-helical domain (Figure 6D). This is referred to as the flex domain structure.

**Figure 6:**
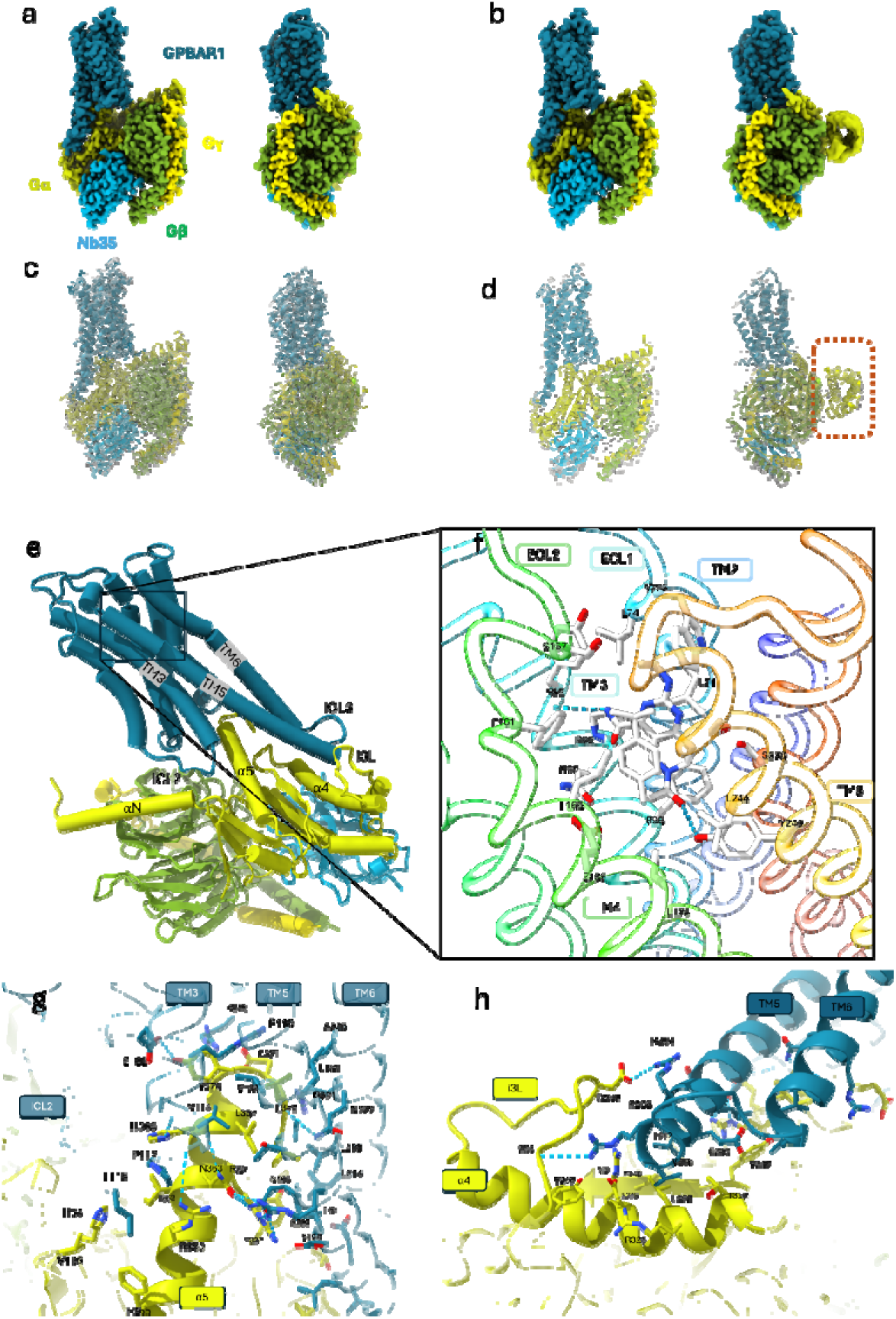
Cryo-EM map and atomic model, (a) Cryo-EM map at 2.5 Å of the GPBAR1 G protein complex, the map has been colored according to the different sub-units. (b) Cryo-EM map at 2.9 Å with the flexible AH domain. (c, d) Atomic model of the two complexes placed in the density. (d) The flexible Gα part is surrounded by dotted lines. (e) Important regions for the interaction between GPBAR1 and Gαs. (f) P395 and the interacting residues of the ligand pocket, the backbones is rainbow colored (Blue N-ter Red C-er). (g) Contact residues between GPBAR1 and the Gαs α5 helix. (h) Contact residues between TM5 and ICL3 of GPBAR1 with α4 and i3L of Ga.

The core structure is used to study the interactions within the complex because it has the highest resolution. GPBAR consists of seven transmembrane domains (TM), three cytosolic loops (ICL), and three extracellular loops. TM3, TM5, and TM6 surround α5 helix of Gαs (Figure 6G), while TM5, TM6, and ICL3 interact with α4 and i3L of Gαs (Figure 6H), incorporating GPBAR into the complex. The high resolution of the structure gives precise positioning of the ligand. The ligand pocket is formed by 16 residues: L71, L74, W75, Y89, P92, N93, F96, S157, F161, L166, E169, Y240, L244, S247, and S270. Most of the contacts are hydrophobic, but a hydrogen bond is formed between the nitrogen of the ligand’s carboxyl amine group and the main chain carboxyl group of Y89. Additionally, the hydroxyl group of Y240 forms a hydrogen bond with the ketone group of P395 (Figure 6F).

GPBAR incorporates into the complex by interacting with the last two helices of Gαs, α5-α4, and the loop i3L (Figure 6G,H). The GPBAR residues involved in the interaction with α5 helix of Gαs are R110, E109, A113, and V114 from TM3; L118 and P117 from ICL2; V188, L189, T191, R194, N195, I199, and D198 from TM5; and L214, L218, R221, N222, and A225 from TM6. The remaining GPBAR contacts are made by the beginning of TM5 and ICL3, where arginine 204 and 208 form hydrogen bonds with D302 and T298 of Gas i3L. Additionally, residues L202, E203, and V206 contact R321, D322, L325, R326, and T329 of α4 helix and Y337 of D6 of Gαs.

The α-helical (AH) domain of the flex domain structure shows the only major difference compared to the core structure. The low resolution of this domain (Supplementary data, Figure B) and the fact that it does not appear in all the classes of the 3D classification step (Supplementary data, Figure C) reflect its high flexibility. This is in line with previous studies showing that the AH domain delocalizes from the GαsRas domain and becomes flexible in the absence of nucleotides[38]. This flexibility is thought to be functionally important, allowing the α-helical domain to undergo essential conformational adjustments for efficient signaling. Note that for the rest of the structure, the organization and contacts between the different sub-units are the same as before.

### 2.4 Complementarity of cryo-EM and HDX-MS to decipher molecular determinants involved in GPBAR1/G complex

To better understand the molecular determinants of interactions between the different biological partners involved in GPBAR1/G complex, the HDX-MS data is finally plotted on the 2.5-Å cryo-EM structure.

Overall, the GPBAR1 HDX-MS data are consistent with the cryo-EM structure of the GPBAR1/G complex (Figure 7). Protection against H/D exchange is also observed for TM7, TM2, TM3, and TM5, in agreement with the orthosteric pocket of the P385 ligand in the GPBAR1 receptor (Figure 7.A). All residues involved in direct interaction with the ligand in the cryo-EM structure are part of the protected peptides identified by HDX. Maximally deuterated control (maxD) experiments (Supplementary data, Figure H) demonstrate that helix 7 behaves differently from the other transmembrane helices. Specifically, for TM7, D uptake profiles are very similar to each other, regardless of incubation times, and are especially close to the maxD values[39] obtained when the protein is denatured. It is, therefore, highly likely that the TM7 helix is unstructured in the Apo form. TM7 helix in GPBAR1 becomes less flexible and forms more contacts with other TM helices upon binding of P385 and Gs proteins (Figure 7B,C). These interactions consequently alter the environment of TM4, TM5, and TM6 helices at the GPBAR1/G protein interface, explaining the differences in incorporation and conformational changes observed.

**Figure 7:**
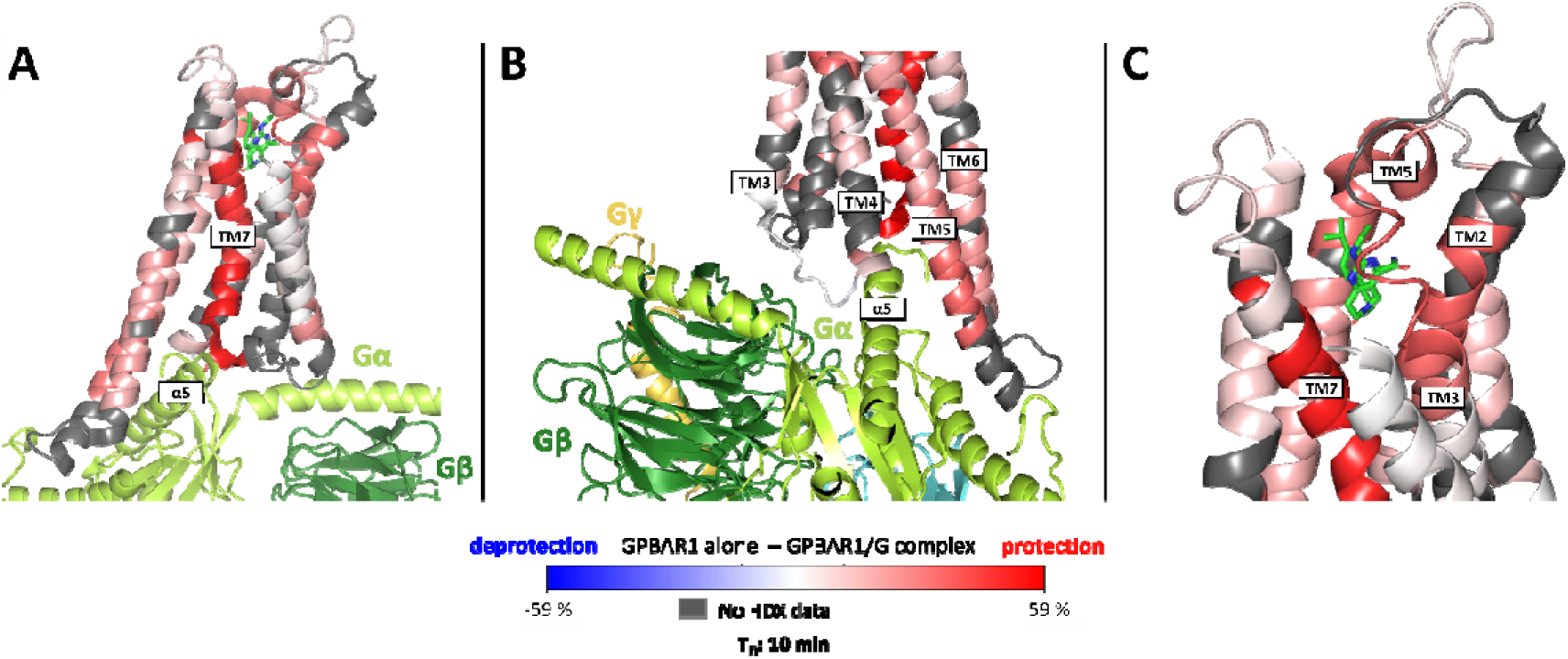
Visualization of D incorporation differences (deuteration time: 10 min) on the GPBAR1 structure. Regions are colored according to the given scale (-59%; 59%); those for which no information is available are colored in gray. (A) Overview of the whole receptor. (B) Zoom on the regions of interaction between the different proteins making up the complex. (C) Zoom on the ligand-binding site.

Additionally, HDX-MS data reveal that while the C-terminal end of the α5 helix is strongly protected (interaction of residues Gln363, His366, Tyr370 and Leu3722 with GPBAR1), the other end of the helix is exposed to the solvent and is therefore not localized inside the receptor. These observations are in agreement with cryo-EM data and general GPCR/G protein binding mechanisms. Cryo-EM data reveal that insertion of the α5 helix compacts the structure of GPBAR1 and further increases conformational constraints in the extracellular domain where the P395 ligand is located. This highly conserved mechanism is characteristic of the interaction of GPCRs with G proteins[31, 32]. In the case of GPBAR receptors, the insertion of helix α5 is not as deep as that of other class A GPCRs, due to the extension of the TM5 and TM6 helices[36, 37].

Furthermore, the cryo-EM structure of the complex enables further interpretation of the HDX data in the context of G-protein subunit interactions (Figure 8). Figure 8A shows the stabilizing effect of Nb35, which makes the region less accessible to the solvent, explaining the protective effect observed on the incorporation profile. On the other hand, the interaction between Gβ region and the αN helix of the Gαs subunit can be depicted in (Figure 8B). The dynamic effect observed above stems from the fact that the αN helix of Gαs progressively “pushes” part of the Gβ subunit. Finally, the structure reveals another interaction interface between Gαs and Gβ subunits, thus explaining the protective effects observed on the α2 helix of Gαs and Gβ. Note that the reduced solvent accessibility for helix α3 of Gαs can also be explained by its interaction with Nb35.

**Figure 8:**
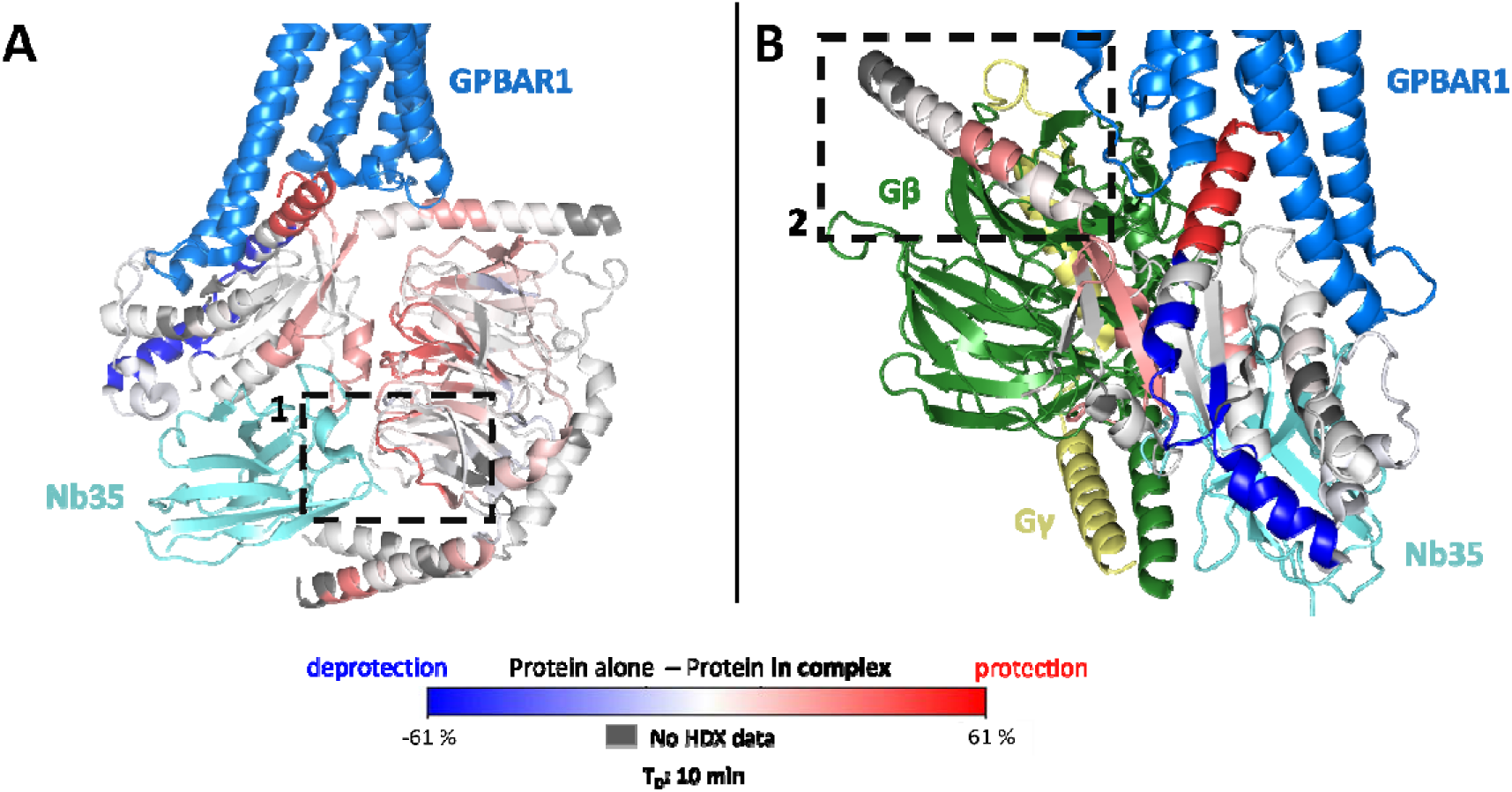
Visualization of D incorporation differences (deuteration time: 10 min) on the G protein structure. Regions are colored according to the given scale (-61%; 61%); those for which no information is available are colored gray. (A) D-incorporation differences are observed on the 3 subunits, Gαs, Gβ, and Gγ. Box 1 shows the direct interaction between Gβ and Nb35. (B) Visualization of incorporation differences on Gαs only. Box 2 shows the interaction between one of the G□s helices and the Gβ subunit.

## 3 DISCUSSION

An exhaustive description of the molecular determinants and the conformational dynamics of active GPBAR1 upon heterodimeric Gs binding obtained by a combination of state-of-the-art mass photometry, HDX-MS, and cryo-EM data is provided. Several technological optimizations were essential to achieve a high-quality dataset for GPCR proteins.

First, to the best of our knowledge, our HDX-MS study is one of the few performed on a fully active full-size GPCR (composed of full-length GPBAR1 bound to its agonist P395 ligand) bound to heterotrimeric Gs (comprising Gαs, Gβ, and Gγ subunits and stabilized by Nb35 nanobody). This leverages the level of complexity of the HDX-MS experiment to a 5-subunit multiprotein complex, which is state-of-the-art for membrane protein analysis. In addition, achieving sequence coverage >60-70% is still challenging for HDX-MS of membrane proteins, especially for GPCRs. Through an extended optimization of the upstream sample preparation, we achieved sequence coverages >99% for GPBAR1, allowing a thorough view of the conformational dynamics throughout the GPCR sequence. Using the latest cryo-EM sample preparation and data collection procedures, we also obtained high-resolution (2.5-Å) structures of the GPBAR1/Gs complex, revealing an additional yet undescribed highly flexible AH region for GPBAR1.

Since these experiments offer a unique opportunity to compare the dynamic insights from HDX-MS with the more static view of cryo-EM, the GPBAR1/Gs study serves as a valuable case study and could be further extended to investigate GPCR/Gs protein systems more broadly. Overall, cryo-EM and HDX-MS are highly consistent and cross-validate each other, highlighting increased local stability of many intracellular and extracellular regions in GPBAR1 when bound to the Gs heterotrimer. This increased stability suggests that G proteins’ binding to the agonist GPBAR1 may reduce the flexibility of specific regions, which could be crucial for receptor activation and signal transduction.

In addition, HDX-MS not only reveal conformational rearrangements on the GPBAR1 side but also on the trimeric Gs. Indeed, HDX-MS shows that the α5 helix of the GDs subunit undergoes a significant decrease in H/D exchange upon complex formation. This finding is in good agreement with cryo-EM data, which reveals a direct interaction of the α5 helix with GPBAR1, suggesting that the conformational changes in GPBAR1 upon activation are transmitted through this interaction to the G protein, facilitating signal transduction.

Finally, the complementarity and synergy of high-resolution (cryo-EM) and lower-resolution (HDX-MS and mass photometry) using a model GPCR/Gs complex is demonstrated. MPhoto provides a rapid way to monitor the formation of GPCR/Gs complexes in a membrane mimic environment. Despite its lower mass accuracy compared to native MS, MPhoto does not require any prior buffer exchange, limiting sample preparation artifacts, especially with membrane proteins. MPhoto affords assessment of membrane protein homogeneity, oligomeric state, or complex formation within a few minutes, offering real-time insights into the assembly process and ensuring the good quality and integrity of the sample prior to more complex and time-consuming experiments like HDX-MS and cryo-EM. The systematic combination of cryo-EM and HDX-MS provide valuable information for the analysis of large multiprotein complexes such as GPCR/G protein complexes. While cryo-EM provides a high-resolution almost static picture of the complex, HDX-MS reveals the dynamic conformational changes that occur during complex assembly and activation. We believe that the systematic integration of such complementary biophysical techniques will provide a more comprehensive understanding of the molecular mechanisms underlying GPCR signaling, which could impact and fasten the development of more targeted and effective therapeutics.

## 4 MATERIALS AND METHODS

### 4.1 Hydrogen Deuterium Exchange – Mass Spectrometry (HDX-MS)

#### 4.1.1 Peptide identification

Before performing HDX-MS experiments, peptides were identified by digesting undeuterated samples using the same protocol and identical liquid chromatographic (LC) gradient as detailed below and performing mass spectrometry experiments in MS^E^ data acquisition mode with a Xevo G2 XS-QTOF mass spectrometer (Waters) over a range of m/z 50–2000. A 0.5 mM sodium formate solution (90/10, 2-propanol/H2O, v/v) was used for calibration, and a 200 pg/µL leucine enkephalin solution (50/50/0.1, H_2_O/ACN/HCOOH, v/v/v) was applied for mass accuracy correction in positive electrospray ionization mode (ESI+) and resolution mode. MS^E^ runs were analyzed using ProteinLynx Global Server (PLGS) 3.0.3 (Waters), and peptides identified in each run, with a minimum intensity of 3000, with at least 0.3 fragments per amino acid, 2 consecutive minimum products and a mass error below 10 ppm were selected in DynamX 3.0 to generate peptides lists (Waters).

#### 4.1.2 Optimization of the HDX conditions

Prior to labeling experiments, an extensive optimization of <u>GPBAR1</u>’ GPCR digestion was carried out. All the tested conditions were performed on <u>GPBAR1</u> samples in duplicates to rapidly evaluate the most efficient protocols and optimal parameters to maximize both sequence coverage and redundancy. The injected quantity of <u>GPBAR1</u> (7, 13 and 26 pmoles), the enzyme used for the online digestion (pepsin and nepenthesin II), the nature of the chaotropic/denaturing agent in the quenching buffer, and the carryover were successively evaluated. Digestions were performed online on a pepsin-immobilized cartridge (Enzymate pepsin column, 300 Å, 5 μm, 2.1 × 30mm, Waters) or a nepenthesin II protease column (2.1 × 20 mm, Affipro, Czech Republic). The best experimental conditions were employed on the other apo biological partners (G-protein, nanobody, and complex) to ensure the quality of their digestion under the same conditions. Subsequently, optimal conditions were used for the total complex. A detailed description of the different buffer compositions tested during the optimization phase is provided in supplementary data.

#### 4.1.3 Labelling experiments

Biological partners – <u>GPBAR1</u> (26 pmol), trimeric G protein (39 pmol) and nanobody Nb35 (79 pmol) – were deuterated at room temperature alone or in the complex. Deuteration was carried out by incubating 2 μL of the stock sample solutions into 36 μL of D2O buffer (20 mM HEPES pD 7.5, 150 mM NaCl, 0.02% LMNG w/v) for different labeling time points (10 s on ice / 10, 60, 600, and 3600 s at RT), corresponding to a 95% deuterated final solution. 38 μL of each deuterated preparation were quenched with 38 μL of quench buffer (4 M urea pH 2.3 and 500 mM TCEP) kept on ice with a final pH of 2.4. Deuteration, quenching, and injection of the samples were manually performed in three technical replicates.

#### 4.1.4 Protein digestion and LC-MS conditions for labeling experiments

Digestion and chromatography were conducted on an Acquity UPLC M-Class System with HDX Technology (Waters, Manchester, UK). Digestion was performed online on a nepenthesin II protease column (2.1 × 20 mm, Affipro, Czech Republic) at a 100 μL/min flow rate of 0.1% formic acid solution at 20 °C for 3 min. Peptides were then trapped on an Acquity BEH C18 VanGuard pre-column (1.7 μm, 2.1 × 5 mm, Waters) and separated on an Acquity UPLC BEH C18 analytical column (1.7 μm, 1.0 × 100 mm, Waters) at 0.1 °C. Peptide separation was performed at a flow rate of 40 μL/min, with an elution gradient of solvent A (0.1% formic acid, water) and solvent B (0.1% formic acid, acetonitrile) from 5 – 35% B over 7 min followed by a 1 min ramp to 85% B. All exchange reactions were performed in triplicate. To eliminate peptide carryover, the protease column was washed three times between two runs of deuterated samples using alternatively 1.5M guanidine-HCl in 500mM glycine buffer (pH 2.3), and a solution composed of 5% acetonitrile, 5% 2-propanol, and 20% acetic acid.

#### 4.1.5 HDX-MS data analysis

After a first round of automated spectral processing using DynamX, isotopic profiles for all identified peptides were manually checked, corrected, and validated. A back-exchange assessment and related correction were performed for each protein following the protocol described by Peterle *et al*[39]. Differences in deuterium uptake were statistically validated with a p-value of 0.01 with the MEMHDX software[40] with statistical significance thresholds.

### 4.2 Cryogenic electron microscopy (cryo-EM)

#### 4.2.1 Grid preparation and data acquisition

4 μL of sample were applied onto an UltrAuFoil grid (Quantifoil R1.2/1.3, 300 mesh), which was rendered hydrophilic by a 90 s treatment in an ELMO glow discharge system operating at 3 mA and in a partial vacuum of 3.6 × 10^-1^ mbar. The grid was then blotted for 10 s at a blot force of 15 and flash-frozen in liquid ethane using a Vitrobot Mark IV (Thermo Fisher Scientific) at 6.5°C and 100% humidity. Images were acquired on a Titan Krios G4 (Thermo Fisher Scientific) operating at 300 kV in nanoprobe mode using SerialEM software version 4.1.0 beta for automated data collection. Movies were recorded on a Falcon 4 direct electron camera after a SelectrisX energy filter using a 10 eV slit width. Images were collected at a magnification of 165,000x (corresponding to a pixel size of 0.731 Å). The defocus range was -0.8 to -1.8 μm. Each movie was composed of 703 frames with a total dose of 50 e/Å^2^. The frames were regrouped into 39 fractions, and the first fraction was excluded during motion correction.

#### 4.2.2 Data processing

All data processing was performed in RELION version 4.0. Figure X provides details of the data processing workflow. The alignment of movie frames, dose weighting, and correction of beam-induced specimen motion were done using RELION’s implementation of MotionCorr. Contrast Transfer Function (CTF) estimation was performed using ctffind4. Picking of particles was carried out using the RELION Gaussian blob picker. Initially, particles were extracted and binned four times to speed up the initial data analysis step. Three rounds of 2D classification were performed to remove images containing damaged or aggregated particles and ice contamination, followed by 3D classification with alignment. The 3D class that best represented our complex, showing high-resolution structural features, was chosen for further analysis and particles were re-extracted at their original size to perform refinement. The obtained alignment was used to perform 3D classifications without alignment, one was with no mask and T=10 to keep the best particles for high-resolution refinement, and the second with a mask and T=50 to classify the flexible region of G alpha. Selected classes from each classification were refined using RELION auto-refine job. Ctf refinement and particle polishing were performed. EM Reayd was used for map sharpening.

#### 4.2.3 Model building

Structure 7CFM was fitted in cryo-EM map and modified using coot version 0.9.8.2. Geometry optimization was performed with servalcat, and final refinement using Refmac servalcat. The map used for refinement was a normal post-process map from RELION.

### 4.3 Protein production

#### 4.3.1 Construction and expression of GPBAR1

The gene for human full-length GPBAR1 (UniProtKB-Q8TDU6) was cloned into pFastBac vector. To facilitate protein expression and subsequent purification, a HA signal peptide followed by a FLAG epitope, a 3C protease site, and thermostabilized apocytochrome b562RIL (BRIL) were added at the N-terminus of GPBAR1. The construct was expressed in Sf21 insect cells using the Bac-to-Bac baculovirus expression system (Thermo Fisher Scientific). The cells were grown in SFM Sf-900 II medium (Gibco) to a density of 4.5×10^6^ cells/ml and infected with the recombinant baculovirus at a multiplicity of infection of 2 at 27 ℃ for 48 hours. Cells were then harvested by centrifugation and stored at -80 °C for future use.

#### 4.3.2 GPBAR1 purification

The cell pellets were thawed and lysed by osmotic shock in 10 mM Tris-HCl pH 8, 1 mM EDTA buffer containing iodoacetamide (2 mg/ml), 1 DM P395 (synthesized by Novalix Tunisia), and complete Protease Inhibitor Cocktail Tablets (Roche). After centrifugation (15 min at 38,000 g), the membranes were solubilized using a glass dounce tissue grinder in a solubilization buffer containing 20 mM Hepes pH 7.5, 100 mM NaCl, 10 DM P395, 1% lauryl maltose neopentyl glycol (LMNG; Anatrace), 0.1% Cholesteryl Hemisuccinate Tris salt (CHS; Anatrace) supplemented with 2 mg/ml iodoacetamide and complete Protease Inhibitor Cocktail Tablets (Roche). The extraction mixture was stirred at 4°C for 1 hour. The cleared supernatant (38, 000 g centrifugation) was loaded by gravity flow onto anti-Flag M2 antibody resin (Sigma-Aldrich). The resin was then washed with 20 column volumes of a wash buffer composed of 20 mM Hepes pH 7.5, 150 mM NaCl, 5 DM P395, 0.06% LMNG and 0.006% CHS. The bound protein was eluted in the wash buffer supplemented with 0.2 mg/ml Flag peptide (Sigma-Aldrich). The eluted protein was concentrated to 500 μl using a 50 kDa spin filter and further purified by size exclusion chromatography on a Superdex 200 Increase 10/300 column (Cytiva) in a buffer containing 20 mM Hepes pH 7.5; 150 mM NaCl, 0.006% LMNG; 0.0006% CHS; 5 DM P395. The fractions corresponding to mostly monodisperse protein were collected, concentrated with a 50 kDa spin filter, and subjected to a second SEC on a Superose 6 Increase 10/300 column (Cytiva) with a buffer containing 20 mM Hepes pH 7.5; 150 mM NaCl, 0.006% LMNG; 0.0006% CHS; 5 DM P395 to reach a better separation from oligomeric or aggregated species. The fractions containing monodisperse protein were pooled and concentrated to 1.4 mg/ml for HDX-MS experiments.

#### 4.3.3 Construction, expression, and purification of G_s_ heterotrimer

Gs heterotrimer was expressed in Sf21 cells grown in SFM Sf-900 II medium (Gibco). Human GDs subunit with 1 to 15 original residues swapped with 1 to 18 residues of GDi (Kawai *et al*., 2020) was cloned in pFastBac vector, while N-terminal 6×His-tagged human GD1, and human GD2 were cloned into pFastBac-Dual vector. The baculoviruses were generated and amplified using the Bac-to-Bac baculovirus expression system (Thermo Fisher Scientific). Sf21 cells cultured in SFM Sf-900 II medium (Gibco), at a density of 4 × 10^6^ cells/ml, were coinfected with both viruses at a 1:1 GDs:GD1D2 ratio for 48 hours at 28°C. Cells were harvested by centrifugation and pellets were stored at −80°C. Cells were resuspended in lysis buffer (10 mM Tris pH 7.4, 100 μM MgCl_2_, 5 mM β-mercaptoethanol (β-ME), 10 μM guanosine diphosphate (GDP,) and complete Protease Inhibitor Cocktail Tablets (Roche)) until pellets thawed. The lysate was spun for 15 min at 38,000 g, and then pellet was homogenized in solubilization buffer (20 mM HEPES pH 7.5, 100 mM NaCl, 1% sodium cholate, 0.05% LMNG, 5 mM MgCl_2_, 5 mM β-ME, 10 μM GDP, 5 mM imidazole, and complete Protease Inhibitor Cocktail Tablets (Roche)) using a glass tissue grinder. Solubilization further proceeded for 1 hour with gentle stirring at 4 °C before insoluble debris was removed by centrifugation at 38,000 g for 30 min. Pre-equilibrated nickel-bound Superflow resin (TaKaRa) was added to solubilized supernatant and the mixture was incubated on rotation device for another 1 hour at 4°C. The Gs-bound resin was collected by centrifugation for 5 min at 4,000 g and transferred to a gravity column. Beads were washed with 10 CV of the wash buffers containing increasing concentrations of LMNG and decreasing concentrations of cholate until a final wash in 50 mM NaCl, 20 mM HEPES pH 7.5, 0.1% LMNG, 1 mM MgCl2, 5 mM β-ME, 100 μM GDP and 20 mM imidazole, before being eluted with the same buffer containing 200 mM imidazole. Then 1 μL Antarctic Phosphatase (NEB) was added and the sample incubated overnight at 4°C. Next day the eluate was diluted two-fold with 50 mM NaCl, 20 mM HEPES pH 7.5, 0.1% LMNG, 1 mM MgCl2, 5 mM β-ME, 100 μM GDP to decrease imidazole concentration, passed through a 0.22-μm filter, and loaded onto a pre-equilibrated Q Sepharose resin. The resin was washed with 10 CV of wash buffer (20 mM HEPES pH 7.5, 100 mM NaCl, 0.02% LMNG, 1 mM MgCl_2_, 100 μM Tris(2-carboxyethyl)phosphine (TCEP), 10 μM GDP), and the bound protein was eluted in the wash buffer containing 250 mM NaCl. The fractions containing Gs heterotrimer were pooled, concentrated to 1-2 ml, and diluted three-fold in 20 mM HEPES (pH 7.5, 0.02% LMNG, 1 mM MgCl_2_, 100 μM TCEP, 10 μM GDP) in a drop-wise manner to set the final concentration of NaCl to 125 mM. Gs heterotrimer concentrated to 11 mg/ml was flash-frozen and stored at -80°C for HDX-MS experiments.

#### 4.3.4 Nb35 expression and purification

Nanobody-35 (Nb35) bearing a C-terminal 6His-tag was expressed in the periplasm of *Escherichia coli* strain BL21 (DE3), and grown in a TB culture medium with 100 μg/mL ampicillin at 37 °C, 200 rpm. When OD600 reached 1, 0.7 mM IPTG was added to induce its expression. Induced cultures were grown overnight at 18 °C. The cells were collected by centrifugation and lysed in 20mM Tris-HCl pH 7,5; 150 mM NaCl; 20% Glycerol; EDTA-free Protease Inhibitor Cocktail tablets (Roche) by sonication. After lysis, cell fragments were removed by centrifugation. The supernatant was collected and incubated for 2 hours with Ni-NTA beads on a rotation device at 4 °C. The beads were collected and washed with a buffer composed of 50 mM Tris pH8, 500 mM NaCl, 20 mM imidazole. The protein was eluted in the wash buffer supplemented with 200 mM imidazole. The eluate was concentrated with 5 kDa spin concentrator and loaded on a Superdex 200 Increase 10/300 column (GE Healthcare) using 20 mM HEPES pH 7.5, and 150 mM NaCl as a buffer. Fractions containing the monodisperse peak of Nb35 were pooled, concentrated with a 5-kDa centrifugal filter concentrator and flash-frozen in liquid nitrogen for later use.

#### 4.3.5 Expression and purification of GPBAR1/Gs complex

Sf21 cells were cultured in SFM Sf-900 II medium (Gibco) at 28 °C to a density of 4.5×106 cells/ml and then were co-infected with <u>GPBAR1</u>, Gαs and Gβ1γ2 using Bac-to-Bac baculovirus expression system (Thermo Fisher Scientific) at the ratio of 1:1:1. The cells were collected by centrifugation 48 h post-infection and stored at -80 °C until use.

Cell pellets from 2 L of culture were resuspended in 30 mM Hepes pH 7.5; 100 mM NaCl; 10 mM MgCl_2_, 0.2 mM TCEP, supplemented with complete Protease Inhibitor Cocktail tablets (Roche). GAPBR/Gs/Nb35 complex formation was initiated by addition of 20 μM P395, Nb35 (10 μg/ml) and apyrase (25 mU/ml, Sigma-Aldrich). The suspension was incubated for 1.5 hours at room temperature and the membrane pellets were collected by centrifugation at 38,000 g for 30 min. The membranes were solubilized using a glass dounce tissue grinder in a solubilization buffer containing 20 mM Hepes pH 7.5, 100 mM NaCl, 10 mM P395, 1% LMNG, 0.1% CHS, and complete Protease Inhibitor Cocktail Tablets (Roche). The sample was further incubated for 1 hour with gentle stirring at 4 °C. The supernatant was collected by centrifugation at 38,000 g for 30 min and then incubated with 5 ml Talon resin for 1 hour at 4 °C. The resin was extensively washed with a wash buffer containing 20 mM Hepes (pH 7.5; 150 mM NaCl, 0.06% LMNG; 0.006% CHS; 0.02% GDN; 2 mM MgCl_2_, 5 μM P395; 100 μM TCEP) and the bound proteins eluted in the same buffer supplemented with 200 mM imidazole. Then the eluate was slowly loaded on anti-Flag M2 antibody affinity resin, washed with a buffer composed of 20 mM Hepes (pH 7.5; 150 mM NaCl, 0.006% LMNG; 0.0006% CHS; 0.02% glyco-diosgenin (GDN; Anatrace); 2 mM MgCl_2_, 5 μM P395; 100 μM TCEP) and eluted in the same buffer supplemented with 0.2 mg/ml Flag peptide. The eluted complex was concentrated and further purified by size-exclusion chromatography using a Superose 6 Increase 10/300 column (GE Healthcare) pre-equilibrated with buffer containing 20 mM Hepes (pH 7.5; 150 mM NaCl, 0.006% LMNG; 0.0006% CHS; 0.02% GDN; 2 mM MgCl_2_, 5 μM P395; 100 μM TCEP). Eluted fractions that consisted of <u>GPBAR1</u>/Gs/Nb35 complex were pooled and concentrated with a 100-kDa centrifugal filter concentrator to 4 mg/ml. The obtained complex was either directly used to prepare cryo-EM grids or flash-frozen in liquid nitrogen for HDX-MS experiments.

## ACKNOWLEDGMENTS

This work was supported by NovAliX, the CNRS, the University of Strasbourg, the “Agence Nationale de la Recherche” and the French Proteomic Infrastructure (ProFI; ANR-10-INBS-08-03). The authors thank Ameur Benyounes & Hajer Abdelkafi for providing P395 compound. Additionally, T.B. thanks Daniele Peterle and John R. Engen for guidance regarding the maxD protocol. J.C. acknowledges ANRT and NovAliX for funding his Ph.D.

## Supplementary material description

### DATA AVAILABILITY STATEMENT

The PDB model and the cryo-EM map have been deposited in the Protein Data Bank and the Electron Microscopy Data Bank, and are accessible using these identifiersPDB: 9GYO, EMD-51700.

The HDX-MS data have been deposited to the ProteomeXchange Consortium via the PRIDE partner repository with the dataset identifiers XD047822, PXD047827 and PXD04783.

