## supplementary data for "Deciphering molecular determinants of GPCR-G protein receptor interactions by complementary integrative structural biology methods"

- **Reproducibility of MPhoto experiments**

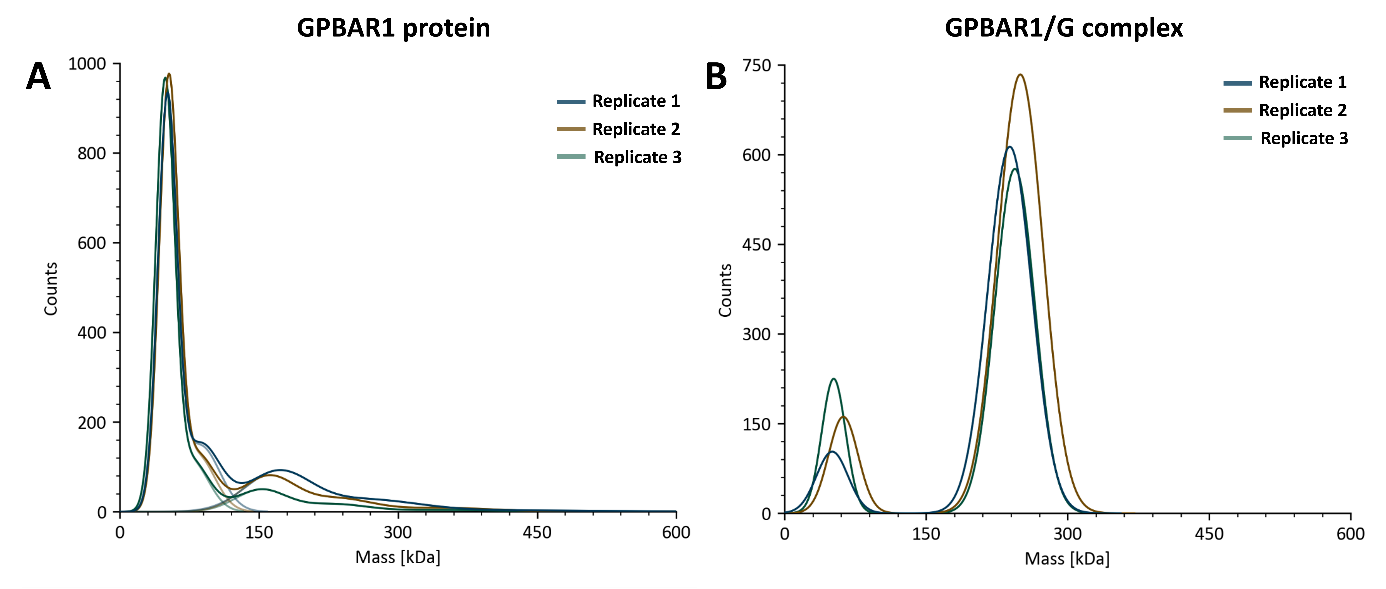

Figure A: Assessment of MPhoto experiment reproducibility. Superposition of analyses of three GPBAR1 (A) and GPBAR1/G protein complex (B) technical replicates. The continuous curve corresponds to the mass distributions of the majority of species fitted with a Gaussian function.

- **CryoEM data processing method**

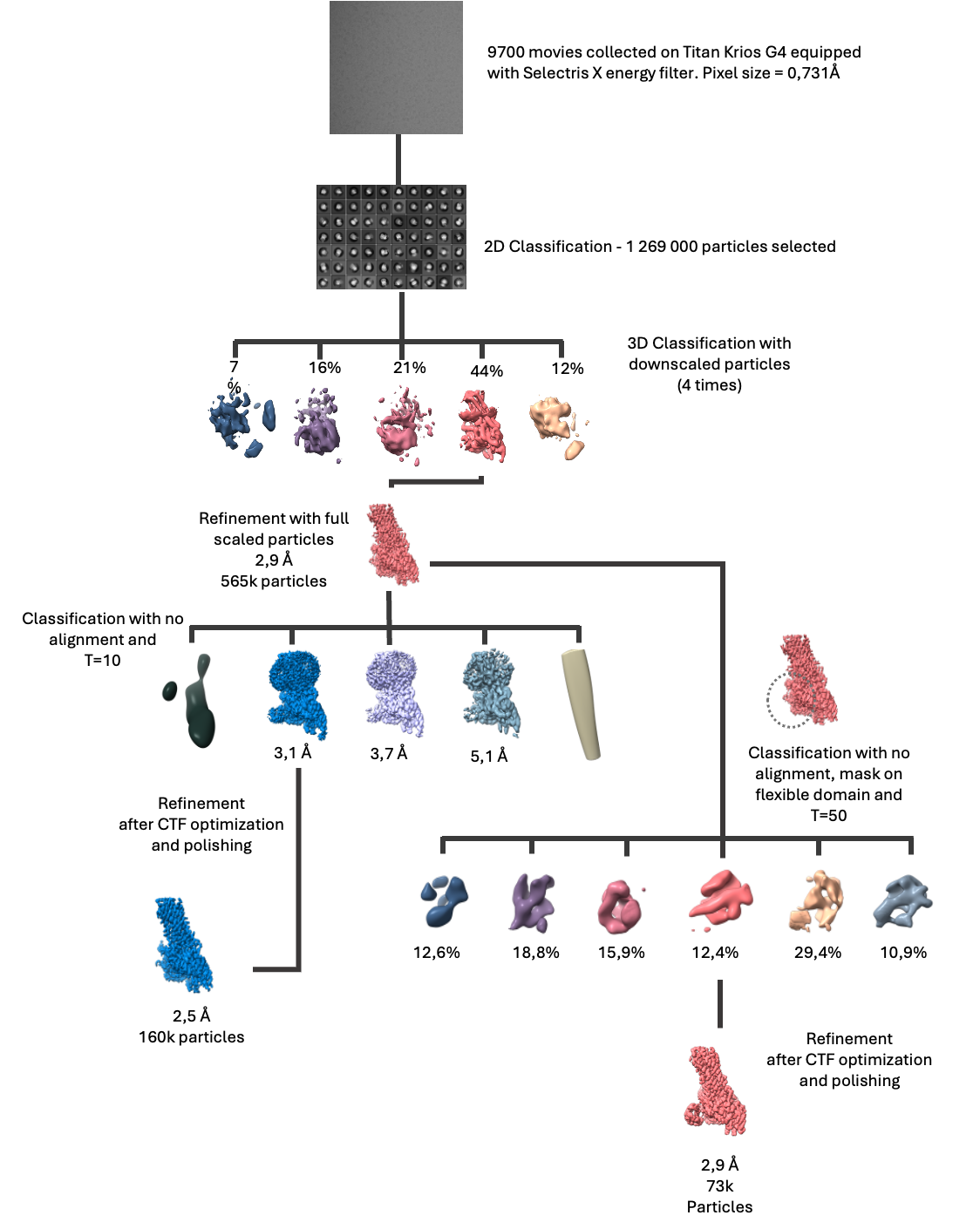

Figure B: Cryo-EM data processing

Schematic diagram of cryo-EM data analysis, showing how cryo-EM maps were obtained. 9700 micrographs were collected. After initial 3D classification, 1,269,000 were selected and classified in 3D. The resulting particles then underwent further 3D classification to refine two high-resolution cryo-EM maps.

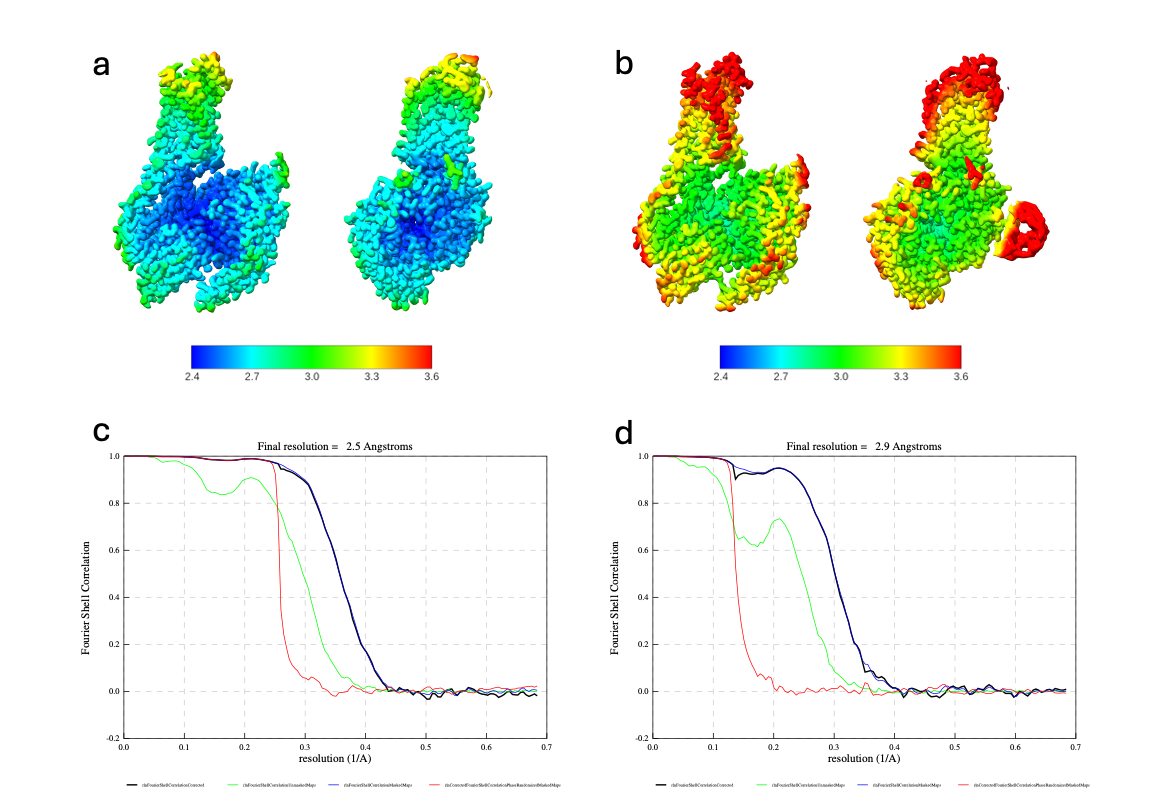

Figure C: Local and global resolution map estimation

(a,b) Local resolution of the two cryo-EM maps obtained, to compare their resolutions, the same color scale has been used. (c,d) FSC curve of the structures obtained.

- **Optimization of protein digestion**

Optimization of the protein digestion was initially carried out on the GPBAR1 receptor alone. Preliminary analyses using standard laboratory conditions showed that optimization was necessary (Figure D). Indeed, while a sequence coverage of 62% may seem satisfactory at first glance, visualization of the peptides identified on the protein sequence indicates that one part of the protein (at the N-terminus) is much better digested than the others. This region, identified in grey in Figure 7, does not correspond to the GPBAR1 receptor but to the fusion protein, heat-stabilized apocytochrome b562RIL (BRIL), used to facilitate receptor production and purification. This protein is soluble and, therefore more easily digested, which explains the observations. The transmembrane domains of GPBAR1, on the other hand, are poorly digested, with only a few peptides identified in these regions.

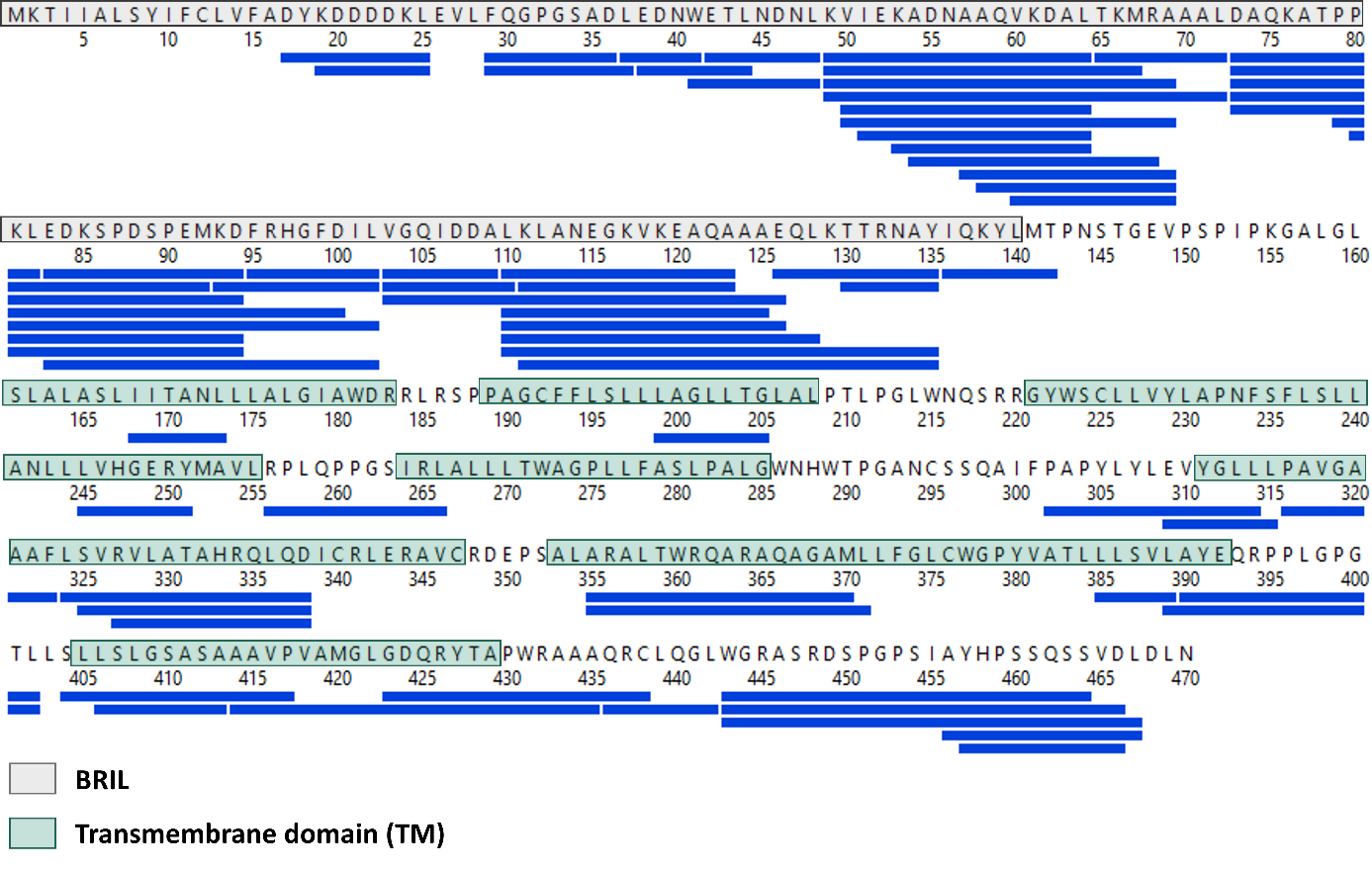

Figure D: Presentation of peptides (blue) identified after pepsin digestion of the sample. 7 pmoles were injected. A Q buffer consisting of 2 M guanidine-HCl was used. Sequence coverage: 61.7%. Number of peptides identified: 70. Redundancy: 3.3.

The following parameters were evaluated: the quantity of protein injected into the spectrometer, the nature of the enzyme used for digestion, the addition of a reducing agent, and the chaotropic agent in the Q buffer used for the HDX-MS experiments.

Optimization of the various experimental parameters during the pre-analytical phase gave excellent results for the digestion of this GPCR, with > 98% sequence coverage and >7 redundancy (Figure E). Although this optimization phase may seem tedious, it remains essential, enabling us in the end to multiply the number of peptides identified by a factor close to 9, and to gain 45% in sequence coverage compared with our initial conditions (7 pmoles injected, pepsin digestion, Q buffer composed of 2M guanidium chloride). What is more, the whole transmembrane domain is now well covered, which is particularly promising for the level of information that HDX-MS experiments will be able to provide.

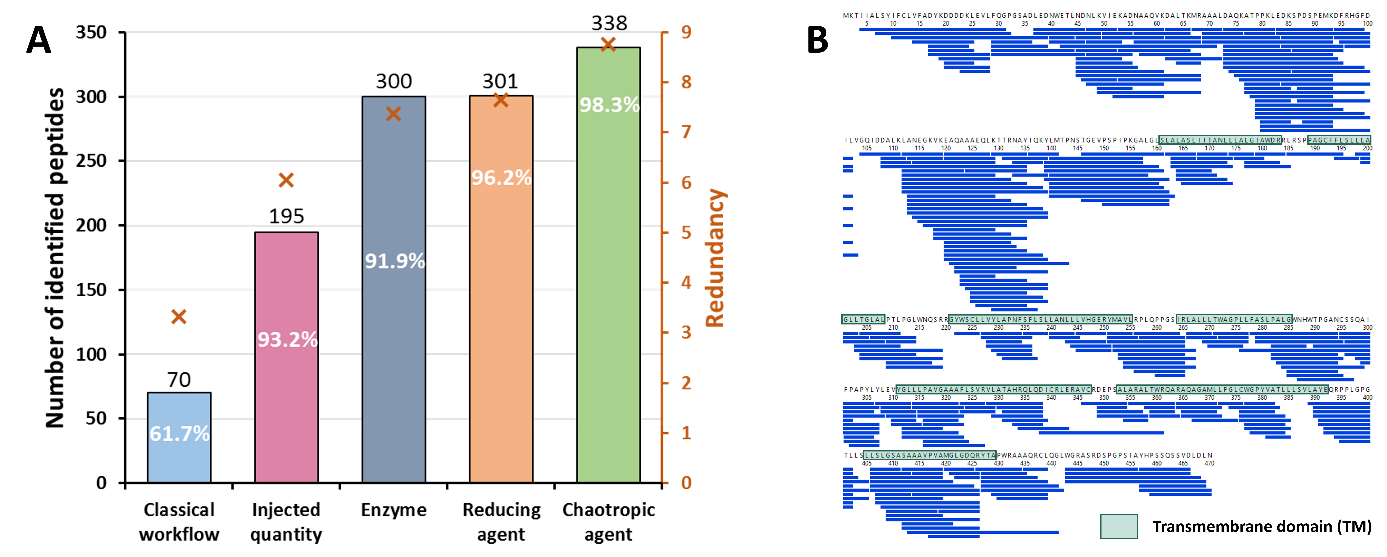

Figure E: Importance of protein digestion optimization for improving the information resolution. (A) Influence of the different optimization steps on the sequence coverage, the number of peptides identified, and the redundancy. (B) Presentation of peptides (blue) identified after nepenthesin II digestion of the sample. 26 pmoles were injected, and a Q buffer consisting of 4 M urea and 500 mM TCEP was used. Sequence coverage: 98.3%. Number of peptides identified: 338. Redundancy: 8.8.

Each of the other proteins, namely the G protein (consisting of its three subunits Gα, Gβ, and Gγ), the nanobody Nb35, and the complex (grouping all partners), were analyzed in duplicate using the same analysis conditions as those for GPBAR1 (Table A). First of all, we can see that the proteins alone and in complex digest very well, with sequence coverages above 90% with very high redundancy values (between 6 and 22). When we look at the results obtained for proteins alone or in complex, a difference can be observed. Indeed, for all proteins, the number of peptides identified is lower when they are analyzed simultaneously within the complex (20% loss of peptides for GPBAR1, for example). This result is consistent with the fact that digestion of a protein complex made up of 5 different subunits generates a very large number of peptides, making it more challenging to identify as many of the peptides associated with each protein (overlap during chromatographic separation, differences in intensity and ionization, etc.). However, the observed reduction in the number of peptides identified still provided very good results, enabling us to carry out HDX-MS experiments on all the biological partners in this study.

Table A: Comparison of sequence coverage, number of identified peptides, and redundancy obtained for GPBAR1, the Gα, Gβ, and Gγ subunits of the G protein and the Nb35 nanobody when the proteins are analyzed alone or in the complex with optimized experimental conditions.

|  | Analyses | GPBAR1 | Gα | Gβ | Gγ | Nb35 |
| --- | --- | --- | --- | --- | --- | --- |
| Protein  alone | **Sequence coverage** (%) | 100 | 99.7 | 100 | 100 | 91.1 |
|  | **Number of peptides** | 219 | 333 | 386 | 96 | 217 |
|  | **Redundancy** | 7.3 | 12.2 | 16.1 | 21.6 | 20.2 |
| Protein  in complex | **Sequence coverage** (%) | 99.4 | 99.7 | 100 | 94,4 | 100 |
|  | **Number of peptides** | 175 | 303 | 315 | 81 | 162 |
|  | **Redundancy** | 6.7 | 11.6 | 13.0 | 18.7 | 16.0 |

- **Nanobody Nb35/G-protein interaction, as a positive control for our HDX-MS experiments**

Before looking more closely at the conformational dynamics of the GPBAR1 receptor during GPCR/G complex formation, we wanted to validate the quality of our dataset. To this end, we sought to confirm the known regions of interaction between the nanobody and the G protein. The results obtained show statistically significant differences in D incorporation over a large part of the Nb35 nanobody (Figure F). The differences observed are positive, meaning that D incorporation of peptides is lower when Nb35 is in complex than when it is alone. These results are consistent with the fact that, when the nanobody interacts with other biological partners, the interaction zones are less accessible to the solvent, which explains the lower D incorporation. What is more, we can see that the strongest differences (above the maximum significance threshold of 10% established by the MEMHDX software) correspond to the 3 CDRs identified for Nb35. These results, therefore, confirm that Nb35 is interacting with the other partners in the complex, involving the expected regions, thus validating our analytical method and giving us confidence in the quality of our dataset and our interpretations.

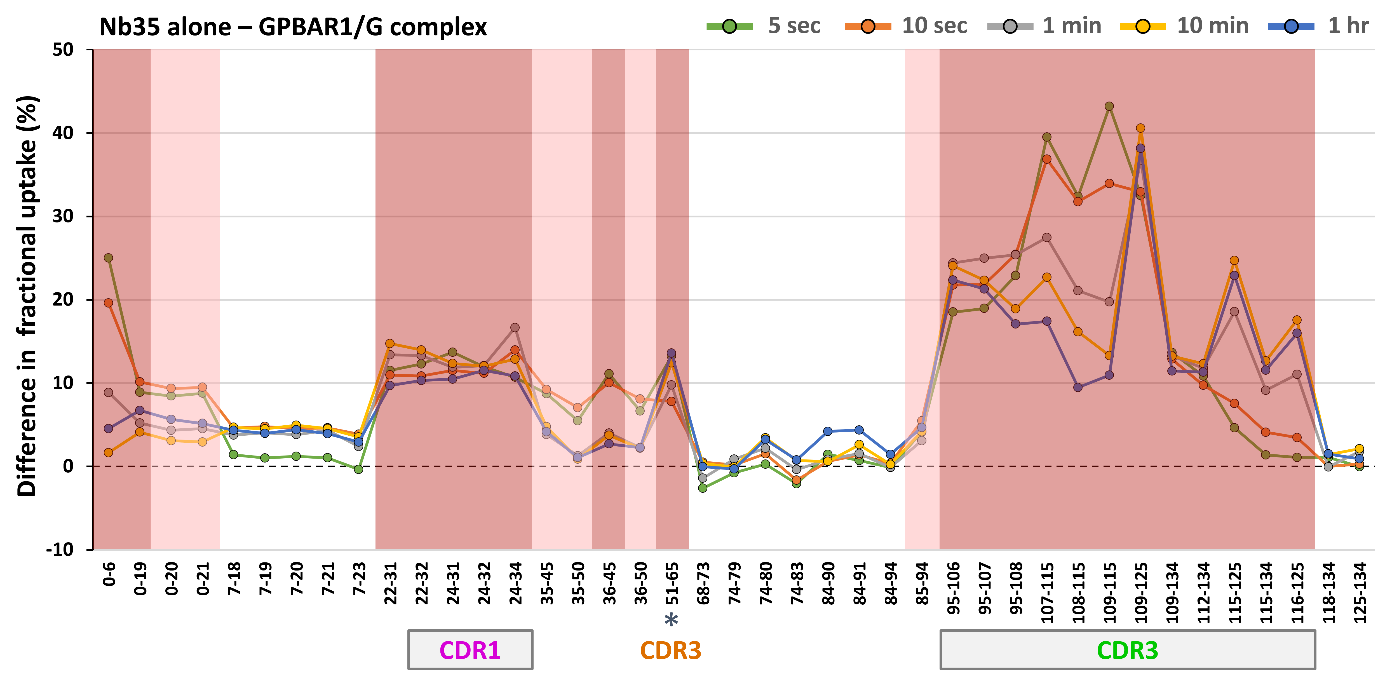

Figure F: Differences in relative D incorporation between Nb35 alone and in complex for each peptide identified at different deuteration times. The framed peptides show statistically significant differences in D incorporation (MEMHDX, p-value 0.01), between 5 and 10% (light red) and above 10% (dark red), showing a decrease in solvent accessibility when Nb35 is in complex.

- **BRIL region, as a negative control for our HDX-MS experiments**

Peptides derived from BRIL protein digestion show no difference in incorporation between the two states (Figure G), confirming the "validity" of the dataset.

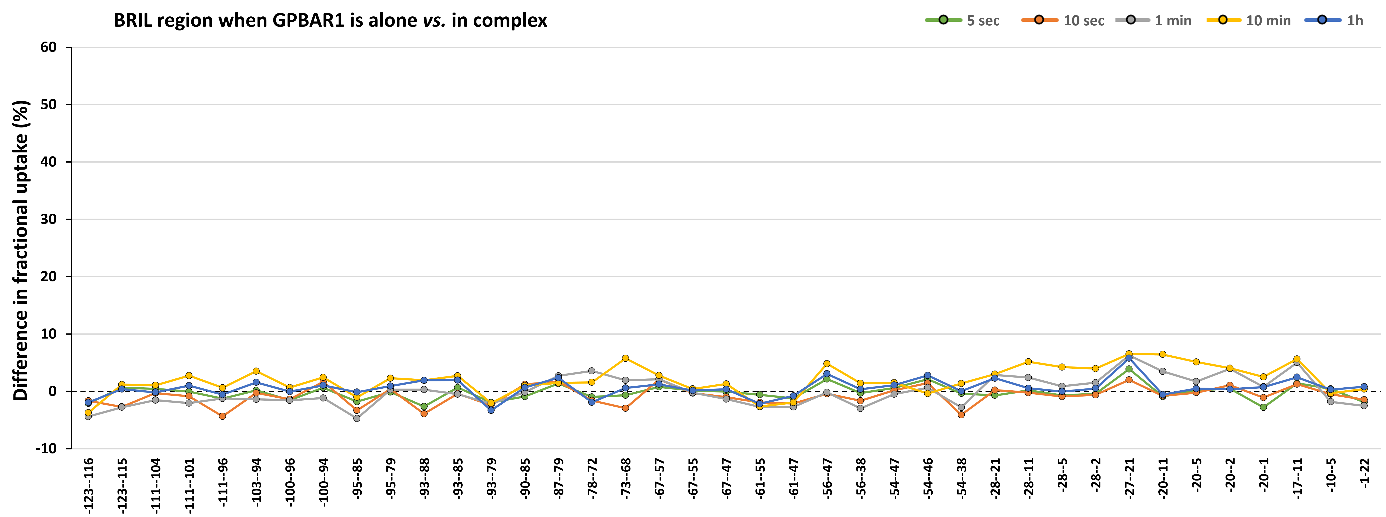
Figure G: Differences in relative D incorporation BRIL region when GPBAR1 is alone and in complex for each peptide identified at different deuteration times. The framed peptides show statistically significant differences in D incorporation (MEMHDX, p-value 0.01).

- **Back-exchange correction**

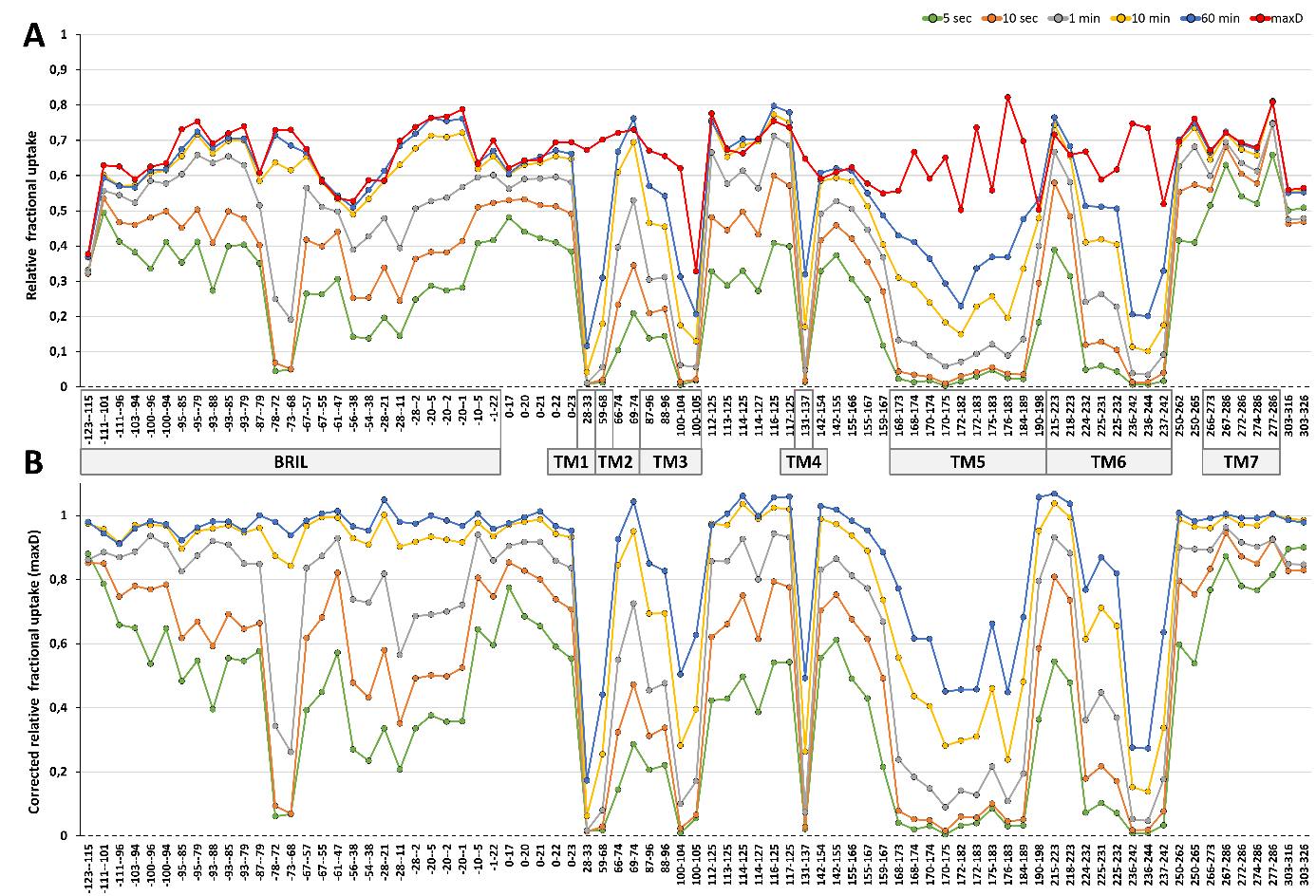

Figure H: Representation of the relative D incorporation of GPBAR1 alone for each peptide identified at different deuteration times without (A) and with (B) back-exchange correction. The peptides resulting from the digestion of the BRIL protein are numbered from -123 to 0 while those resulting from GPBAR1digestion are numbered from 1 to 326.
